# Long-term all-optical interrogation of cortical neurons in awake-behaving non-human primates

**DOI:** 10.1101/274308

**Authors:** Niansheng Ju, Rundong Jiang, Stephen L. Macknik, Susana Martinez-Conde, Shiming Tang

## Abstract

Whereas optogenetic techniques have proven successful in their ability to manipulate neuronal populations in species ranging from insects to rodents—with high spatial and temporal fidelity—significant obstacles remain in their application to non-human primates (NHPs). Robust optogenetics-activated behavior and long-term noninvasive monitoring of target neurons, have been challenging in NHPs. Here we present a method for all-optical interrogation (AOI), integrating optical stimulation and simultaneous two-photon imaging of neuronal populations in the primary visual cortex (V1) of awake rhesus macaques. A red-shifted channel-rhodopsin transgene (C1V1) and genetically-encoded calcium indicators (GCaMP5 or GCaMP6s) were delivered by AAVs, and subsequently expressed in V1 neuronal populations for months with high stability. We achieved optogenetic stimulation using both single-photon (1P) activation of neuronal populations and two-photon (2P) activation of single-cells, while simultaneously recording 2P calcium imaging in awake monkeys. Optogenetic manipulations of V1 neuronal populations produced reliable artificial visual percepts. Together, our advances show the feasibility of precise and stable all-optical interrogation of cortical neurons in awake NHPs, which may facilitate broad applications in high-level cognition and pre-clinical testing studies.

## INTRODUCTION

Optogenetic techniques enable the functional characterization of neuronal populations and circuits with high spatial and temporal precision, but have been understudied in NHPs^1-7^. The successful application of optogenetics to NHPs is critical to discovering mechanistic pathways unique to primates, including: visual processing circuits found only in humans and primates (the only mammals with retinal foveas)^8,9^; high-level cognition circuits^10-15^; and aberrant mechanisms underlying human neurological and psychiatric disorders^16-18^. NHP studies are also essential for pre-clinical testing of optogenetic therapies before they can be translated to human application translated for human use^13,19,20^.

Previous studies have attempted to use traditional electrophysiological techniques to record optogenetic activation. Yet, repeated electrode recordings of the same neurons are difficult to achieve in NHPs. In addition, examining opsin expression patterns online within the area targeted by viral vector infusions, while maintaining the health of the neurons, is not currently possible without 2P laser-scanning microscopy^16,21^. These combined hurdles call for an AOI approach to the application of optogenetic methods in NHPs.

AOI is achieved by the combination of optogenetics to perturb neuronal activity, and calcium or voltage indicators for non-invasive simultaneous readout.^22-27^ AOI’s implementation thus allows monitoring large neuronal populations repeatedly and non-invasively, at single-cell resolution,^28-30^ while enabling detailed mapping of neural circuits during behavior.^31,32^ Pioneering efforts to apply AOI in NHPs combined optogenetics with both *in vivo* epifluorescence imaging and intrinsic signal optical imaging.^19^ Whereas these techniques allowed for large-field viewing, the spatial resolution of the readout was limited, and specific neurons of interest could not be interrogated repeatedly across recording sessions.

Here we combined wide-field 1P and single-cell 2P optogenetic stimulation techniques with recently developed 2P imaging technique in awake macaques^4,33^ to achieve AOI in NHPs. A red-shifted opsin C1V1 and calcium indicators GCaMP5G/GCaMP6s were delivered into V1 by adeno-associated virus (AAV) and expressed in V1 neuronal populations. The labeled V1 neurons exhibited stable repeatable responses to either optogenetic or visual stimulation. The behavioral experiments confirmed that robust artificial visual perception could be induced by optogenetic stimulation of V1 neuronal populations.

## RESULTS

### Transgene Delivery and Expression

We infected area V1 neurons in three monkeys with C1V1 (AAV9-CamKIIα-C1V1(T/T)-ts-EYFP)—a red-shifted channel-rhodopsin transgene—and GCaMP5G/GCaMP6s (AAV1-hSyn-GCamP5G / AAV1-Syn-GCamP6s)—calcium indicators of activity. Six weeks after virus injection, a 1cm-diameter round optical window (glass coverslip attached to a titanium ring) was implanted onto the cortical surface using dental acrylic cement attached to the bone surrounding the craniotomy.^33^

We imaged layer II/III neurons in the infected cortical area using 2P (Fig. 1a). Dark cell bodies indicate C1V1-ts-EYFP expression localized to the membrane^34^ (see also Supplementary Fig. 1). Fluorescence of GCaMP6s was relatively weak in the absence of cellular responses to either visual or optogenetic stimulation.

**Fig. 1.**
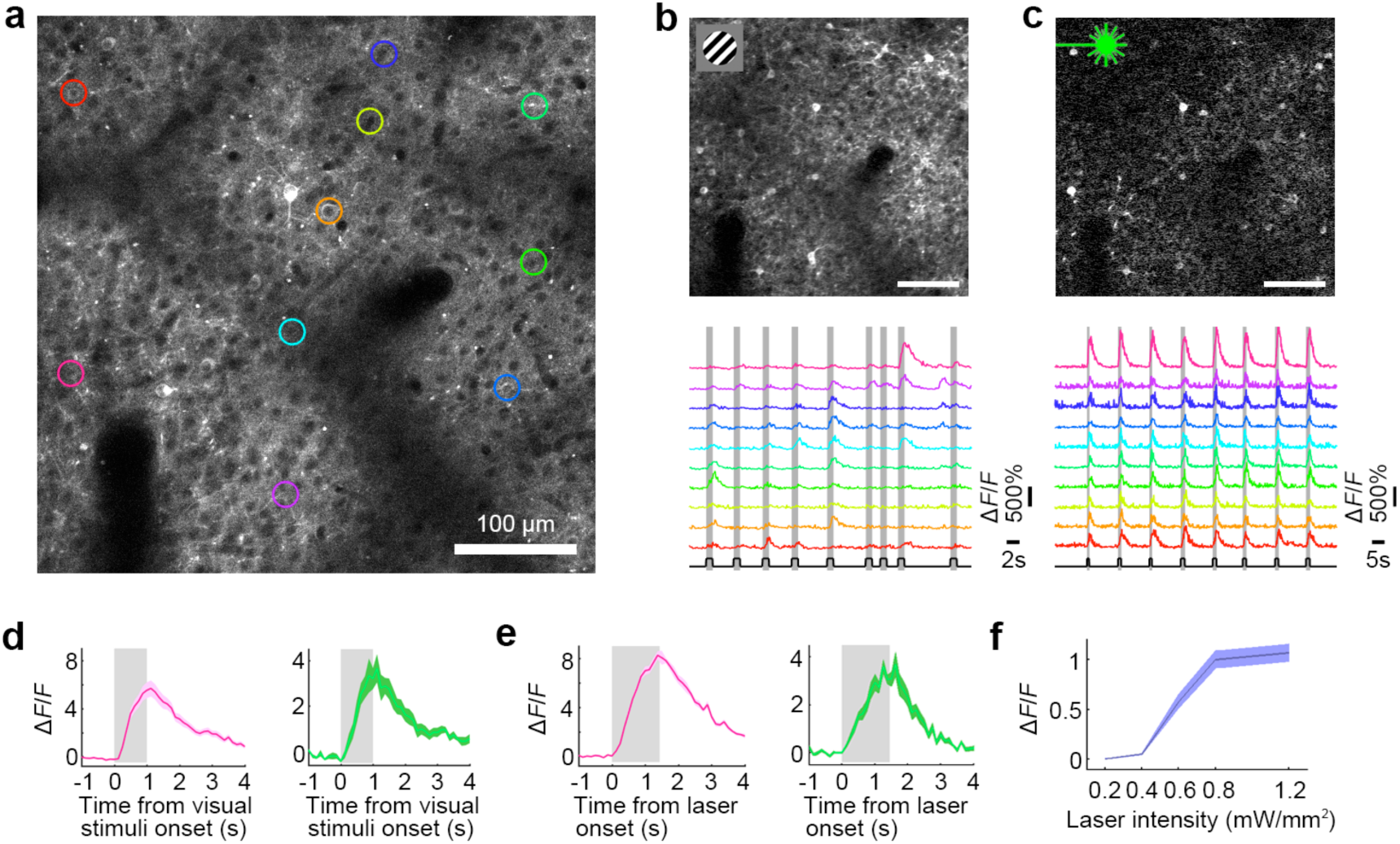
All-Optical Interrogation (AOI) of a V1 neuronal population in awake macaque. **a**, 2P image of V1 neurons expressing C1V1-ts-EYFP and GCaMP6s. The colored ROIs indicate neurons that responded to both visual and optical stimuli, targeted for further analysis. **b**, Top, a differential image of GCaMP6s fluorescence (stimulated-baseline (F-F0), averaged across all stimulation frames), driven by visual stimuli consisting of gratings and color patches. Bottom, calcium signals from 10 neurons (colors from panel **a**) in response to 9 varied visual stimuli (presentation times in gray). **c**, Top, widefield optogenetic stimulation (0.8 mW/mm^2^, 30 Hz and 25% duty ratio) evoked robust responses in the same neurons. Bottom, 8 sequential identical optogenetic stimulations evoked equivalent responses in each cycle. **d**, Responses of two neurons to their preferred visual stimuli (pink, color patch; green, drifting grating; mean ± s.e.m., n = 10 trials). **e,** Optogenetically stimulated responses of the same two neurons are comparable to their visual responses (in panel **d**; mean ± s.e.m., n = 9 trials). **f**, Optogenetic dose-response curve (mean ± s.e.m., n = 9 trials). Optogenetic stimulation saturates at approximately 0.8 mW/mm^2^.

### Simultaneous Optical Manipulation and Two-Photon Readout

NHPs kept eye fixation, while visual stimuli consisting of drifting gratings and color patches were presented sequentially on the neuronal receptive field for 1 second at a time, with > 2 second inter-stimuli intervals. We recorded robust neuronal calcium responses that showed normal orientation and color selectivity, as well as spatial organization (Fig. 1b and Supplementary Fig. 2 and 3).

**Fig. 2.**
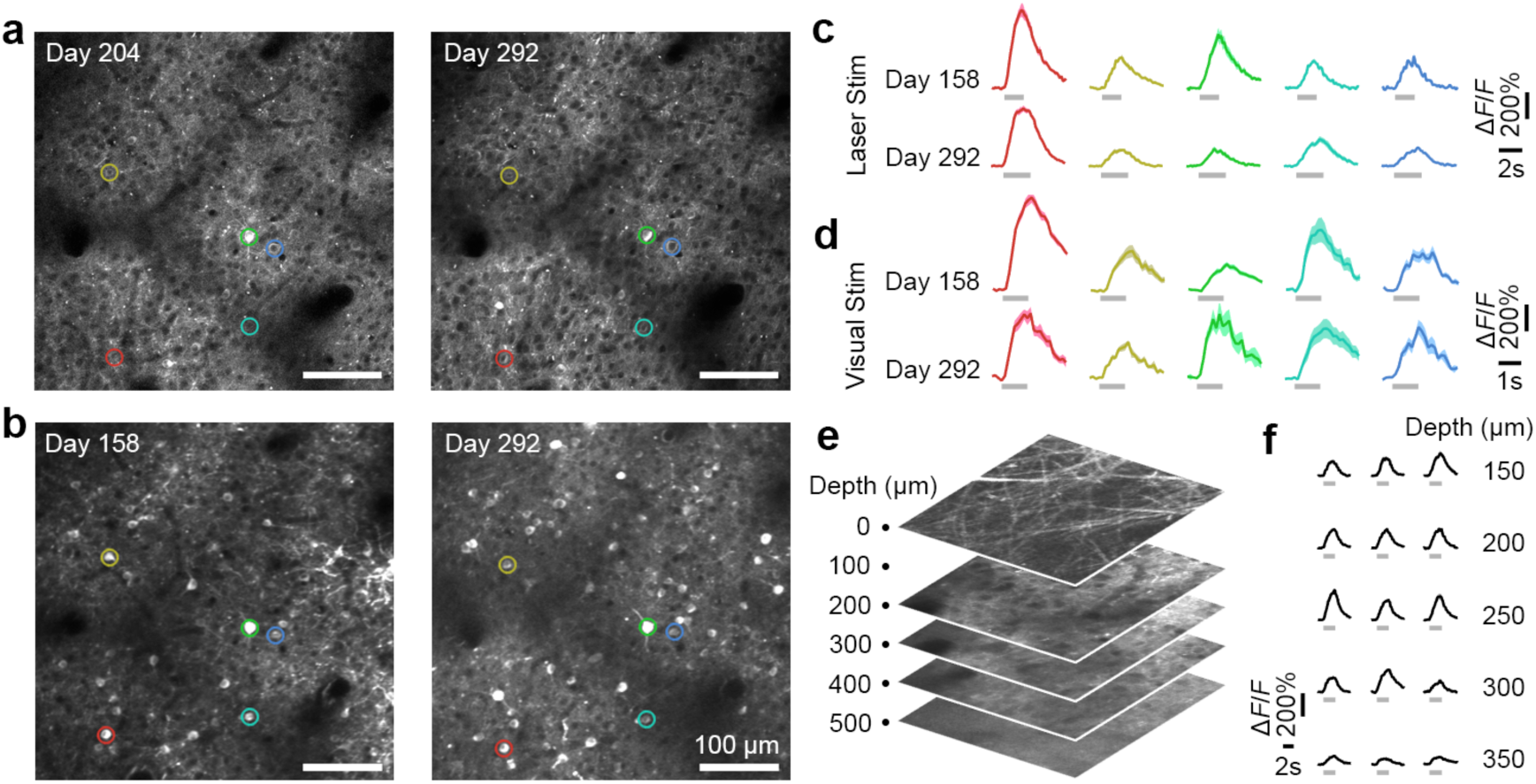
Long-term stability of AOI in macaque V1. **a**, 2P images of a V1 neuronal population on Day 204 and Day 292 after virus injection. **b**, The same neuronal population was activated with visual stimuli on Day 158 and Day 292. **c**, Repeated optical stimulation evoked stable neuronal activity in 5 example cells (colored circles in **a-b**) on Day 158 and Day 292 (mean ± s.e.m., n = 9 trials). **d**, Visual stimulation also evoked repeatable neuronal activity in the same 5 cells (from **a-c**) on Day 158 and Day 292 (mean ± s.e.m., n = 10 and 5 trials respectively). **e**, Multi-layer 2P volume from the cortical surface to 500 µm depth. **f**, Optogenetic calcium responses in different layers (five layers from 150 µm to 350 µm depth, three neurons from each layer) under 1P stimulation.

**Fig. 3.**
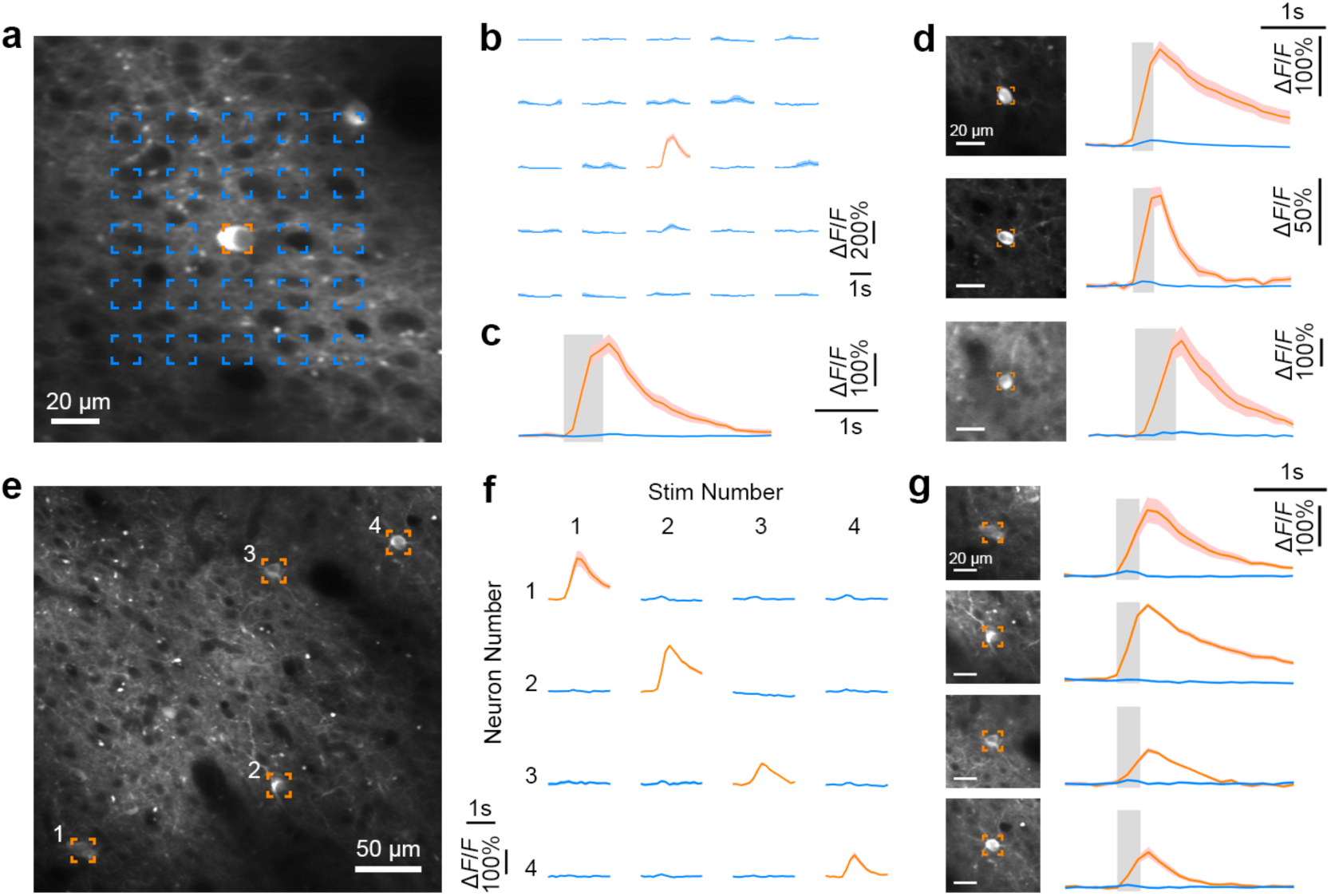
AOI with single-cell resolution in V1. **a**, 2P image of a targeted neuron co-expressing GCaMP5G and C1V1 (orange box). **b**, Calcium responses of the targeted neuron following spiral 2P optogenetic stimulation at each of the 5*5 grid locations from **a**. The target neuron responded only to directly focused 2P stimulation, indicating that 2P stimulation is spatially precise. **c**, Average fluorescence traces from the neuron when targeting two-photon stimulation directly at the soma (orange) versus the surrounding parenchyma (blue; mean ± s.e.m., n = 15 trials). **d**, Fluorescence traces of three other neurons under 2P stimulation. **e**, 2P image containing four targeted neurons co-expressing GCaMP5G and C1V1. **f**, Calcium responses of the targeted neurons (from **e**) when they were sequentially activated with spiral 2P stimulation. Each row demonstrates one of four neurons’ activity (numbered as in **e**) during stimulation on either the same or alternate numbered neuron (columns). **g**, 2P images of the targeted neurons (**e**) and their individual average fluorescence traces when each of them were targeted (orange) versus not targeted (blue; mean ± s.e.m., n = 10 trials).

We then stimulated the neurons optogenetically. Using single-photon stimulation (532 nm laser), we illuminated the entire imaged field (a 1 mm^2^ laser spot), while measuring neuronal activity with simultaneous 2P imaging. Simultaneous optical stimulation and two-photon imaging present a significant challenge: this is because, though the stimulation and recording wavelengths may be sufficiently separated and filtered optically, the optogenetic stimulation power is orders of magnitude higher than the fluorescence power emitted by the activated cells. Thus, stimulation light leaks through the filters, and into the highly amplified photomultipliers (PMTs), with higher power than the relatively small GCaMP fluorescence signal. To address this issue, we shuttered the stimulation path while 2P-scanning the central 75% of the field. This allowed us to view the central area artifact-free (Supplementary Fig. 4).

**Fig. 4.**
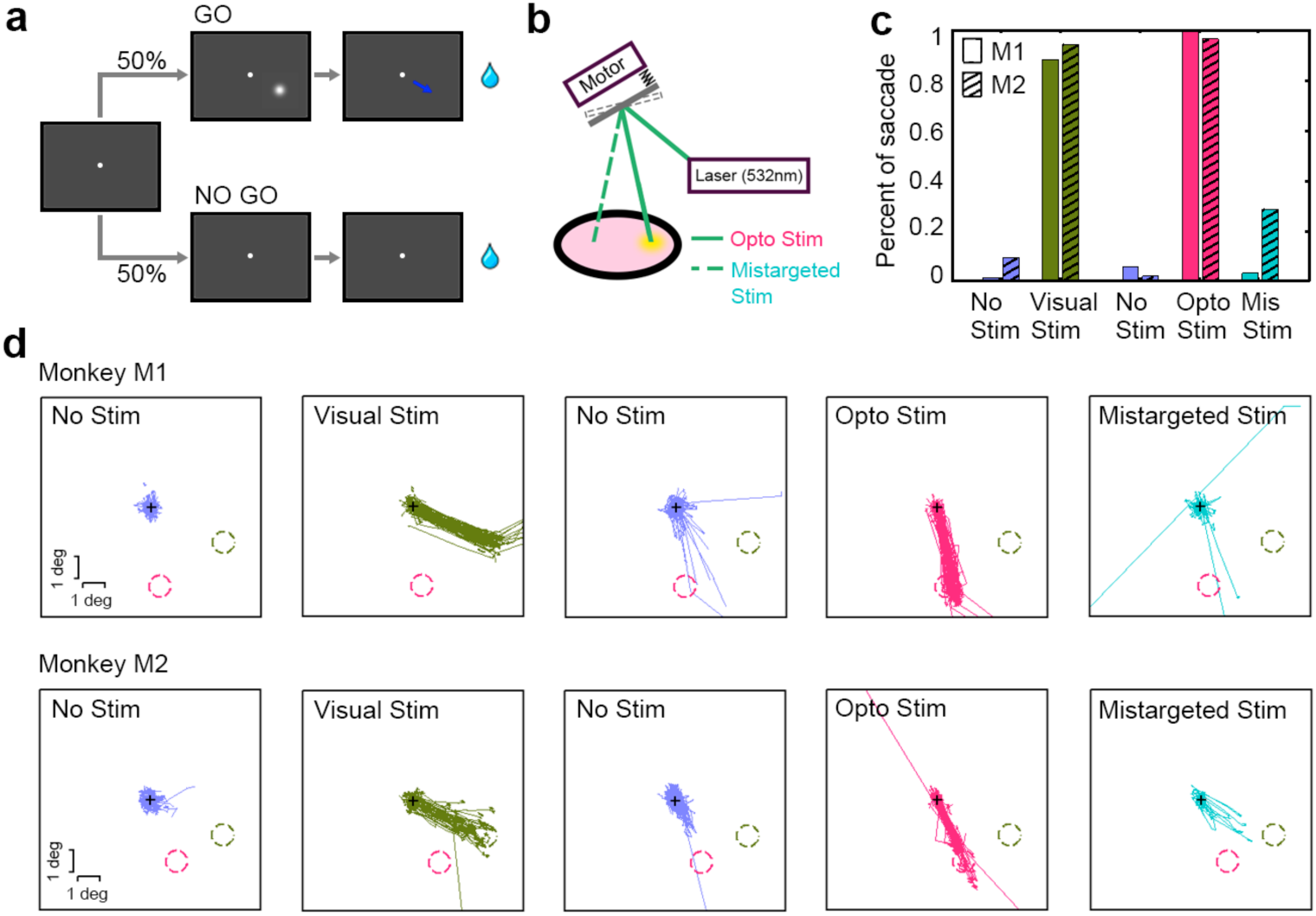
Behavior induced by visual versus optogenetic stimulation in V1. **a,** GO/NO GO visual object detecting task. Monkeys were trained to report the onset of a visual cue (either a visual stimulus—Visual Stim—or an artificial visual perception induced by optogenetic stimulus—Opto Stim) by producing an eye movement. Each trial began when the NHP fixated the central fixation point. In the GO condition, monkeys were required to make a saccade within 500 ms of the cue onset, to obtain a juice reward. In the NO GO condition, no stimulus was presented, and monkeys were tasked with maintaining fixation for 2000 ms, to get a juice reward. **b**, An apparatus used for redirecting laser pulses towards a nearby cortical area not transduced with C1V1 (Mistargeted Stim, see insert). **c**, Analysis of behavior in two monkeys (M1 & M2) during the Visual Stim block (No Stim (blue), and Visual Stim (green)) versus the Opto Stim block (No Stim (blue), Opto Stim (red) and Mistargeted Stim (teal)).**d,** Saccadic trajectories under each condition. The green dash circles indicate the location of the Visual Stim, and the red dash circles denote the receptive fields of C1V1-expressing sites. In Opto Stim, both monkeys uniformly targeted their eye movements to the receptive fields of the C1V1-expressing site.

The optogenetically stimulated cells constituted a considerable fraction of the targeted population, and presented vigorous responses (Fig. 1c). By repeatedly conducting both optogenetic and visual stimulation we made three observations: 1) responses from the two modes of stimulation were comparable to each other in both amplitude and dynamics; 2) repeated stimulation resulted in similarly-sized responses; 3) optogenetic activation did not alter the receptive field properties of neurons that were subsequently stimulated with visual stimuli (Fig. 1d,e). Notably, the dose-response curve revealed that the average laser-evoked responses were saturated at approximately 0.8mW/mm^2^, indicating high sensitivity of the optical manipulation system (Fig. 1f).

### Assessment of Long-Term Stability

Using AOI, we assessed the long-term stability of transgene expression and function in behaving monkeys. Transgene expression level and pattern were maintained (Fig. 2a), and neurons exhibited stable responses and tuning to visual stimuli (Fig. 2b,d and Supplementary Fig. 2 and 3), over several months. The same neuronal population was repeatedly and stably activated by optical stimulation over a four-month period (Fig. 2c). We also evaluated the transgene expression at different cortical depths, from the surface to 500 µm. At 10-months post-infection there was abundant expression between 150 to 300 µm (Fig. 2e), and neurons in this range of depths responded robustly to optical stimulation (Fig. 2f).

Thus, both expression level and neuronal function remained stable over long time periods (in our experience, six months or more) in NHP cortex.

### All-Optical Single-Cell Resolution Stimulation and Recording

A powerful way to assess neural circuit function—for example, to determine the precise weights of the connections between identified neurons in an ensemble—is to photostimulate an individual neuron (without stimulating unwanted targets) while simultaneously monitoring the activity of the connected neurons in the network^5,18,32^. To perform simultaneous single-cell resolution 2P optogenetic activation with 2P calcium imaging of the neuronal population, we added a second optical path to our microscope—with a mode-locked femtosecond laser (λ = 1070 nm, 50 fs) and applied 2P stimulation—and used spirally scanning galvanometers to stimulate the soma of the target cell^31^.

To examine the spatial specificity of 2P activation, we measured the calcium response of the targeted neuron as a function of multiple surrounding stimulation sites (5 x 5 grid) (Fig. 3a). We sequentially stimulated each of the grid’s sites using 2P spiral activation. Robust responses were evoked only when the target neurons soma was specifically targeted (Fig. 3b-d), suggesting that high-spatial precision of spiral 2P stimulation must be focused on the neuronal cell body for strong optogenetic activation to occur. We then simultaneously monitored and sequentially manipulated several neurons in one imaging field (Fig. 3e). Each of these neurons generated strong responses only when targeted by the 2P activation laser (Fig. 3f,g).

### Optogenetic Manipulation of Behavior

To examine the monkeys’ perceptual experience resulting from optogenetic stimulation of V1 neuronal populations, we designed an GO/NO GO visual object detection task, in which two monkeys were required to report the appearance of a visual cue using eye movements (Fig. 4a).

Each trial began when the NHP fixated the central fixation point. Subsequently, a 0.5-degree Gaussian white dot was presented for 22 ms at an eccentricity of approximately 3 degrees as a visual cue for GO (an eye fixation break), and the NHP was rewarded for producing a saccade within 500 ms. On the NO GO trials (50%, no visual cue), the animal was rewarded for holding fixation for 2000 ms for the entire trial. Training proceeded until the NHPs conducted this task with high accuracy (> 80% correct rate; Visual Stim; Fig. 4c). Notably, both monkeys tended to make eye movements towards the location of the visual cues (Fig. 4d, green), though any saccade exceeding one degree in magnitude was sufficient to receive a reward.

We then examined the artificial visual perception generated by optogenetic stimulation—Opto Stim. The GO condition here had no visual cue. Instead, we conducted optogenetic stimulation (a 532 nm laser, 22 ms pulse, subtending 1 mm^2^) at the position of the C1V1-expressing cortex (which was about 3 degrees eccentric from the fovea, in a different position from the stimulus in the Visual Stim block). Similar to the Visual Stim condition, monkeys in Opto Stim received a juice reward if they produced a saccade (> 2 degrees) after the optogenetic stimulation. Both monkeys performed this task well in Opto Stim condition with 99% and 96% accuracy in monkey M1 and monkey M2, respectively (Opto Stim; Fig. 4c). The eye movements correctly targeted the visual space locations of the RFs corresponding to the C1V1 expressing cortical sites (which were never otherwise targeted with Visual Stim cues). This further confirmed that optogenetic stimulation successfully induced artificial visual perception in the NHPs (Opto Stim; Fig. 4d).

To rule out the possibility that any of the observed effects were due to artifacts resulting from the physical side effects of the laser illumination, we also interleaved Mistargeted Stim trials (8.3%) in the GO condition: here, we redirected the laser to a region of V1 cortex that did not express channel-rhodopsin (Fig. 4b). This mistargeted laser should not be capable of evoking optogenetic activation of neurons, or the ensuing artificial visual perception. This control condition was treated as a GO task, and monkeys were again rewarded only for saccades launched immediately after laser onset (< 500ms). Despite the incentive to produce saccades in any direction, we observed significantly fewer saccades in this condition (p < 10^−20^ for M1 and p < 10^−10^ for M2), (Opto Stim versus Mistargeted Stim; Fig. 4c), indicating that the monkeys truly were not aware of the mistargeted laser stimulation.

We also studied saccadic latencies as a function of stimulus type and duration. For Visual Stim, saccadic responses were swift and robust (Fig. 5a), and exhibited consistent latencies of ∼119 ms, measured as the time between cue onset and the saccade crossing the 1-degree magnitude threshold (Fig. 5f). During Opto Stim (2.4 mW/mm^2^), we found that laser pulses of 44 ms duration (or more) elicited robust responses (Fig. 5b-d), and that optogenetic-induced saccadic latencies were ∼30 ms shorter, averaging 90 ms after laser onset (Fig. 5g and Supplementary Fig. 5). This ∼30 ms difference presumably arises from the optogenetic stimulation bypassing the subcortical visual pathway, an observation consistent with previous studies of visual signal propagation from the retina to V1^35,36^.

**Fig. 5.**
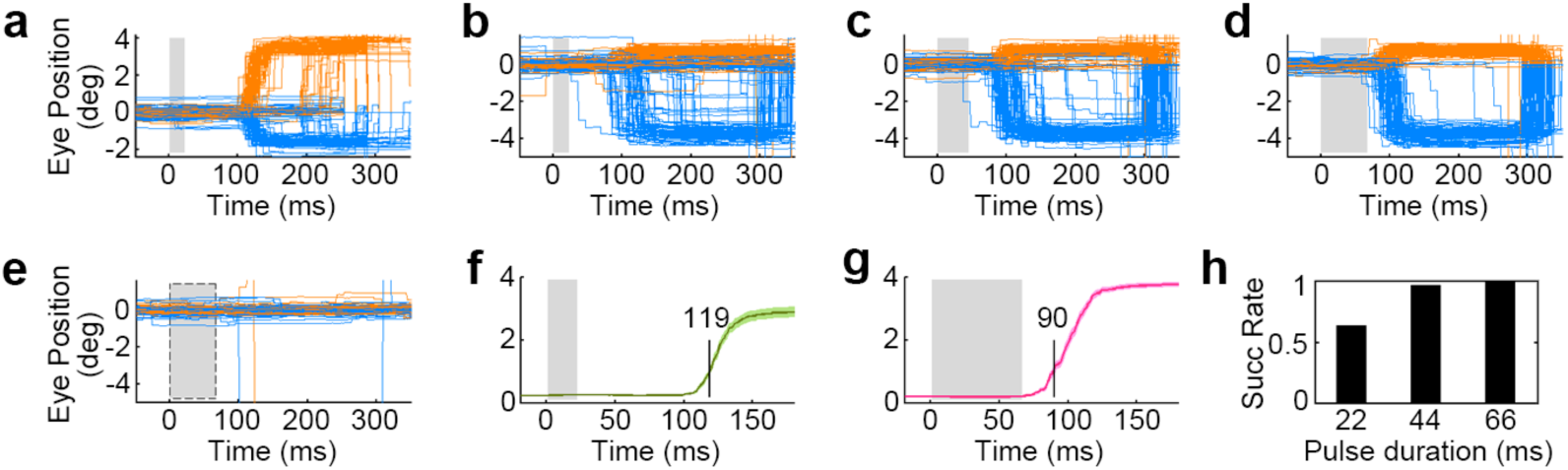
Latencies of saccades triggered by optogenetic activation vs visual stimulation. **a,** Saccades triggered by visual stimuli (single flash of 22 ms). **b**-**d,** Saccades triggered by different optogenetic stimulation pulse modulation durations (22, 44 and 66 ms) on the target (C1V1 expressing) cortical area of monkey M1. Orange and blue lines represent horizontal and vertical eye positions, respectively. **e,** Eye traces when the stimulation laser (66 ms pulse) was targeted to an unlabeled region of cortex. **f**-**g,** Average saccade distance to the fixation point in response to visual (**f**) versus optogenetic (**g**) stimulation (the gray area denotes the stimulation period; mean ± s.e.m., n = 94 and 96 trials respectively). Visually-induced saccades (green line in **f**) crossed the 1-degree threshold with an average latency of 119 ms, whereas laser-induced saccades (red line in **g**) crossed the 1-degree threshold with an average latency of 90 ms. **h,** Success rates of reactions under Opto Stim with varying laser pulse durations.

## DISCUSSION

Optogenetic applications in NHPs have facilitated our understanding of sensory processing, decision making, and the bases of cognition^11,12,16,21,37^, and will likely play a key role in future brain-computer-interfaces, neural prosthetics, and methods to counteract cognitive decline in the aging human brain. As such, optogenetic techniques are undergoing rapid translation to human clinical use. A critical step in the approval, implementation and efficacy of optogenetic therapies will be pre-clinical testing in NHPs, for which methods are currently lacking.

AOI is a particularly powerful optogenetic tool, as it can precisely map NHP neural circuits. Here we have combined optogenetic stimulation with 2P calcium imaging of neuronal responses, to achieve AOI in awake behaving macaque monkeys, in which we co-infected V1 neurons with C1V1 and GCaMP6s, and monitored calcium signal using 2P microscope while stimulating optogenetically. Our experiments revealed robust repeated neuronal responses to both optical stimuli and optogenetic activations, which remained high in quality for many months (Fig. 1,2). 2P optogenetic stimulation moreover evoked strong neuronal responses with single-cell resolution (Fig. 3). Optogenetic milliwatt-level stimulation in V1 cells produced strong and specific responses in identified visual cells. Finally, we compared optogenetically-derived to visually-derived perception by assessing the dynamics of saccadic eye movements produced in response to both modes of stimulation (Fig. 4,5). Together, the above results demonstrate the high sensitivity and stability of our AOI strategy.

### Expression of Optogenetic Actuator and Calcium Indicator in Monkey Cortex

Channelrhodopsin-2 (ChR2) is a commonly used optogenetic actuator for NHPs, though it often requires high laser power to evoke neuronal and behavioral responses^14-17,21^. The high conductance and red-shifted absorption spectrum of C1V1 makes it a preferable choice^12,34,38,39^. This is especially true for AOI experiments, since C1V1‘s excitation spectrum is well separated from that of GCaMPs^31,32^. Expression of C1V1-ts-EYFP was robust, and we observed membrane-localized EYFP fluorescence, which has previously indicated membrane localization of C1V1^38^ (Fig. 1 and Supplementary Fig. 1). We visualized GCaMP6s fluorescence and filtered the widefield 1P stimulation pulses with a 500 ± 12.5 nm filter to block most of the EYFP fluorescence. Although the imaging quality was somewhat reduced due to the filter, we nevertheless identified robust responses derived from both visual and optical stimulation (Fig. 1,2). Note that co-expression of C1V1-ts-EYFP and GCaMP6s in single neurons did not occur with high efficiency, and many neurons were activated by wide-field illumination but not by single-cell photostimulation. We conclude that more powerful molecular tools and gene delivery techniques further advance their utility in NHPs.

### Noninvasive AOI of Neuronal Population in Behaving Monkeys

Two-photon calcium imaging enables the monitoring and tracking of large neuronal populations at single-cellular resolution, over long periods^33,40-42^. AOI methods provide the means to manipulate neural ensembles—or individual neurons—to identify their specific properties and to study their roles in local circuits^31,32^.

AOI using C1V1 moreover decreases tissue damage caused by repeated probe penetration, and the photodamage effects that occur with the ChR2 construct^18,20^. A primary limitation of our method arises from the 2P imaging-depth limit. This confines our AOI to superficial cortical circuits, lying within 500 µm of the surface^43^. New multiphoton microscopy methods will improve and extend the depth limit of AOI to as deep as 1 mm^44^. Cellular-resolution imaging of subcortical structures is currently achievable with fiber-optic confocal laser endomicroscopy (CLE) techniques^45,46^.

### Behavior Effects Induced by Optical Stimulation

Electrical microstimulation of the visual cortex evokes phosphene perception in humans, as well as saccadic eye movements in NHPs^36,47-50^. Similarly, optical stimulation of monkey V1 has been reported to induce saccades^15^, as also observed here. One refinement of our current design, as compared to prior work, is the inclusion of a control condition that targeted an unlabeled region of cortex, to rule out potential non-optogenetic artifacts related to laser activation. By requiring NHPs to make eye movements within a short period of time, we moreover verified that the animals made behavioral choices that directly derived from the optogenetic stimulation of V1 neuronal ensembles with receptive fields in specific regions of visual space, as compared to choices driven by visual stimulation.

### Application and Perspective

Because of the tight homology between the human brain and the NHP brain, the functional characterization of neurons and neural circuits underlying high-level cognition—and cognitive decline—as well as neurological and psychiatric disorders, remains heavily dependent on NHP research. Primates moreover are the only foveate mammals—having unique visual capabilities and oculomotor behaviors^8,9^—which makes NHPs a critical animal model for human visual perception and the development of visual prosthetics. By integrating optogenetics and calcium imaging, AOI offers the ability to precisely determine and manipulate fine functional maps in real-time during NHP behavior. One of AOI’s main functions is the precise manipulation of single neurons and simultaneous monitoring of connected neuronal activity, to determine the strength of connectivity within neural circuits, without unwanted activation of nearby targets.

## METHODS

Detailed methods are provided in the online version of this paper.

## ACKNOWLEDGMENTS

We thank Peking University Laboratory Animal Center for excellent animal care. We acknowledge the Janelia Farm program for providing the GCaMP5-G construct, specifically Loren L. Looger, Jasper Akerboom, Douglas S. Kim, and the Genetically Encoded Calcium Indicator (GECI) project at Janelia Farm Research Campus Howard Hughes Medical Institute. This work was supported by the National Natural Science Foundation of China No. 31730109, National Basic Research Program of China (2017YFA0105201), National Natural Science Foundation of China Outstanding Young Researcher Award 30525016, a project 985 grant of Peking University, Beijing Municipal Commission of Science and Technology under contract No. Z151100000915070, and a U.S. National Science Foundation grant to SLM and SMC (1734887).

## AUTHOR CONTRIBUTIONS

S.T. N.J. conceived the project and designed the experiments. N.J. and R.J. performed experiments. S.T., N.J. and R.J. analyzed the data. S.T., N.J., R.J., S.M.C. and S.L.M. interpreted results and wrote the paper.

## ONLINE METHODS

Rhesus monkeys (*Macaca mulatta*) were purchased from Beijing Prima Biotech Inc and housed at Peking University Laboratory Animal Center. The study used three healthy adult male monkeys, 4 – 6 years of age and weighing 5 – 7 kg. All experimental protocols were approved by the Peking University Animal Care and Use Committee.

### Surgery procedures and implantation of the optical window

Two sequential sterile surgeries were performed on each animal under general anesthesia. In the first surgery, a 16-mm diameter craniotomy was created in the skull over V1. We opened the dura and injected 200 nl of a 1:1 mixture of AAV1.Syn.GCaMP6s.WPRE.SV40 (CS0564, titer 2.2e13 (GC/ml), Penn Vector Core) or AAV1.hSyn.GCaMP5G.WPRE.SV40 (V4102MI-R, titer 2.2e13 (GC/ml), Penn Vector Core) and AAV9.CamKIIa.C1V1.TS.eYFP.WPRE.hGH (V4545MI-R, titer 1.6e13 (GC/ml), Penn Vector Core). After the injections, we sutured the dura, replaced the skull cap with titanium screws, and closed the scalp. The animal then returned to its cage for recovery, and was administered Ceftriaxone sodium antibiotic (Youcare Pharmaceutical Group Co. Ltd., China) for one week. A second surgery was performed 45 days later to implant the head-posts and imaging window^33^.

### Behavioral task

We trained each monkey to sit in a primate chair with its head restrained while performing visual fixation and behavioral choice tasks. Eye position was monitored with an infrared eye-tracking system (ISCAN, Inc.) at 120 Hz. Each trial started with eye fixation on a small white point (0.1^°^, within a window of 1^°^). Visual stimuli were generated using the ViSaGe system (Cambridge Research Systems) and displayed on a 17” LCD monitor (Acer V173, 80Hz refresh rate) positioned 45 cm from the animal’s eyes. Receptive fields of C1V1 and GCaMP expressing sites were localized with small patches of drifting oriented gratings.

We designed a GO/NO GO detection task for monkeys, in which they reported on either Visual Stim or Opto Stim cues by making saccades (Fig. 4a), in two blocks.

In the first block, the Visual Stim block, a real visual object was presented on the monitor as a GO cue. This visual object was flashed for 22 ms at a distance of ∼3 degrees from the fixation point. The NHPs were trained to generate a saccade immediately (within 500 ms) after cue onset, to obtain a juice reward. The central fixation point remained unchanged for the duration of the trial.

In the Opto Stim block, a 1P laser pulse (with a wavelength of 532nm, 1.0 mm diameter, 0.2-2.4 mW/mm, a duration of 22, 44 or 66 ms) was projected onto the C1V1-expressing cortical site in each monkey as a GO cue instead of a real visual object. We interleaved trials with either mistargeted laser stimulation of the cortex (to an area without C1V1 expression; 8.3% trials; Mistargeted Stim in Fig. 4b-d) or without laser stimulation (66.7% trials; No Stim in Fig. 4c-d), as control trials, allowing us to rule out artifacts related to laser operation.

### Two-photon imaging

After a recovery period of 10 days from the second surgery,the animals were trained to fixate their gaze on a fixation point. Two-photon imaging was performed using a Prairie Ultima IV 2P microscope (Bruker Nano, Inc., FMBU, formerly Prairie Technologies) and a Ti: Sapphire laser (Mai Tai eHP, Spectra Physics) with a 16× objective (0.8-N.A., Nikon). The wavelength of the laser was set at 1000 nm for two-photon imaging of GCaMP. Fast resonant scanning (up to 32 frames per second) was used to obtain images of neuronal activity (8 fps by averaging every 4 frames).

### Single-photon optical stimulation

A 532 nm laser was used for 1P optical stimulation. The laser was directly pointed at the target cortical area through the imaging window. Due to the brightness of the stimulation laser and the high-sensitivity of the photomultipliers, a 500 nm band pass (25 nm width) filter was inserted before PMT of green channel during simultaneous 2P imaging. Nevertheless, simultaneous stimulation light could have potentially leaked through the filtering system to cause recording artifacts. We addressed this potential confound by blocking the 532 nm laser light during the scanning of the central image during 2P recordings, using an electronic circuit that shuttered the laser pulse at central 75% of the viewing frame along the Y-axis of each 2P frame (Supplementary Fig. 3).

### Single-cell two-photon activation

A secondary femtosecond laser with 1070 nm wavelength (maximal power, 2.3 watts; pulse width, 100 fs; Fidelity, Coherent, USA) was used on a secondary galvanometer path in the 2P microscope (Ultima IV, Prairie,Bruker, USA) to perform 2P optogenetic activation targeting single cells, while simultaneously recording calcium activity. Spiral regions (5 rotations, 1.2 expansion rate, 0.01 pixel/µs and 30 repetitions) were defined to point target photo-activation areas (Supplementary Fig. 6). The laser power was adjusted to 30 mW at the end of the objective with a polarization beam splitter into 1070 nm femtosecond laser light pathway.

### Image data processing

All data analyses were performed using customized Matlab software (The MathWorks, Natick, MA). The images from each session were first realigned to a template image (the average image of 1000 frames in the middle of an imaging session) using a normalized cross-correlation-based translation algorithm, to correct the X-Y offset of images caused by the motion between the objective and the cortex.

### Strategy for randomization and/or stratification

The order of visual stimuli was randomized during experiments.

### Inclusion and Exclusion Criteria of any Data

No data were excluded during analysis.

### QUANTIFICATION AND STATISTICAL ANALYSIS

Statistics were performed using customized Matlab software. Data are presented as mean ± SEM or as individual data points, as stated in the figure legends. Number of replicates can be found within the figure legends for each experiment.

## DATA AND SOFTWARE AVAILABILITY

The relevant data and corresponding codes can be found in https://github.com/EastRainju/Opto-TP.

## ADDITIONAL RESOURCES

The genetic constructs used in this work are available via Addgene.

